# A *RAD9*-dependent cell cycle arrest in response to unresolved recombination intermediates in *Saccharomyces cerevisiae*

**DOI:** 10.1101/736850

**Authors:** Hardeep Kaur, GN Krishnaprasad, Michael Lichten

**Affiliations:** Laboratory of Biochemistry and Molecular Biology, Center for Cancer Research, National Cancer Institute, Bethesda, Maryland 20892

**Author notes:** Department of Biochemistry and Structural Biology, University of Texas Health Science Center San Antonio, Texas 78229 (Hardeep Kaur). Correspondence: Michael Lichten, Building 37 Room 6124, 37 Convent Dr MSC4260, NIH, Bethesda, MD 20892-4260.

**Keywords:** double Holliday junction dissolution, Sgs1-Top3-Rmi1 helicase-decatenase, DNA damage checkpoint, mitotic cell cycle, homologous recombination

## Abstract

In *Saccharomyces cerevisiae*, the conserved Sgs1-Top3-Rmi1 helicase-decatenase regulates homologous recombination by limiting accumulation of recombination intermediates that are precursors of crossovers. *In vitro* studies have suggested that the dissolution of double-Holliday junction joint molecules by Sgs1-driven convergent junction migration and Top3-Rmi1 mediated strand decatenation could be responsible for this. To ask if dissolution occurs *in vivo*, we conditionally depleted Sgs1 and/or Rmi1 during return to growth, a procedure where recombination intermediates formed during meiosis are resolved when cells resume the mitotic cell cycle. Sgs1 depletion during return to growth delayed joint molecule resolution, but ultimately most were resolved and cells divided normally. In contrast, Rmi1 depletion resulted in delayed and incomplete joint molecule resolution, and most cells did not divide. *rad9Δ* mutation restored cell division in Rmi1-depleted cells, indicating that the DNA damage checkpoint caused this cell cycle arrest. Restored cell division in *rad9Δ*, Rmi1-depleted cells frequently produced anucleate cells, consistent with the suggestion that persistent recombination intermediates prevented chromosome segregation. Our findings indicate that Sgs1-Top3-Rmi1 acts *in vivo*, as it does *in vitro*, to promote recombination intermediate resolution by dissolution. They also indicate that, in the absence of Top3-Rmi1 activity, unresolved recombination intermediates persist and activate the DNA damage response, which is usually thought to be activated by much earlier DNA damage-associated lesions.

## Introduction

The conserved STR/BTR complex, composed of the RecQ-family helicase Sgs1 (BLM in many organisms), topoisomerase III (Top3, Top3*α* in mammals), and RecQ-mediated genome instability protein 1 (Rmi1, BLAP75 in humans), has important functions that maintain genome integrity (Bernstein *et al*. 2010; Larsen and Hickson 2013; Crickard and Greene 2019). STR complex components have two principal biochemical activities: Sgs1/BLM is a 3’ to 5’ helicase that unwinds DNA (Bennett *et al*. 1998); reviewed in (Chu and Hickson 2009; Bernstein *et al*. 2010); and Top3-Rmi1 has robust single strand DNA passage but weak supercoil relaxing activities (Cejka *et al*. 2012). *In vitro* studies have revealed STR/BTR activities that can both promote and limit homologous recombination. Sgs1 acts with the Dna2 nuclease to catalyze DNA end-resection, producing 3’-ended single strand DNA that can invade homologous duplex sequences to initiate homologous recombination; this activity is stimulated by Top3-Rmi1 (Zhu *et al*. 2008; Gravel *et al*. 2008; Cejka *et al*. 2010; reviewed in Mimitou and Symington 2009). Other STR/BTR activities have the potential to act later, to limit the formation of crossover (CO) recombinants (Figure 1). STR/BTR disassembles model D-loop structures, analogs of initial strand invasion products (van Brabant *et al*. 2000; Bachrati *et al*. 2006; Fasching *et al*. 2015). This prevents formation of double Holliday junction joint molecules (dHJ-JMs) that are potential CO precursors (Szostak *et al*. 1983), and directs events towards a process called synthesis-dependent strand annealing (SDSA) that produces noncrossover (NCO) recombinants (Gloor *et al*. 1991). STR/BTR also has an *in vitro* activity called dissolution, in which helicase-driven convergent HJ migration is coupled with Top3-Rmi1-catalyzed strand passage, to take apart dHJ-JMs and produce NCOs (Wu and Hickson 2003; Wu *et al*. 2006; Plank *et al*. 2006). These two activities can have different consequences. Since D-loop disassembly takes apart an early intermediate with the same number of strand breaks as the original lesion, it recreates a lesion that can undergo additional rounds of invasion and disassembly before it is repaired, and thus can redirect events to different recombination pathways (De Muyt *et al*. 2012; Kaur *et al*. 2015; Piazza and Heyer 2019). In contrast, dHJ dissolution acts on an intermediate in which all initial strand breaks have been repaired, producing a mature NCO and thus terminating the recombination process.

**Figure 1.**
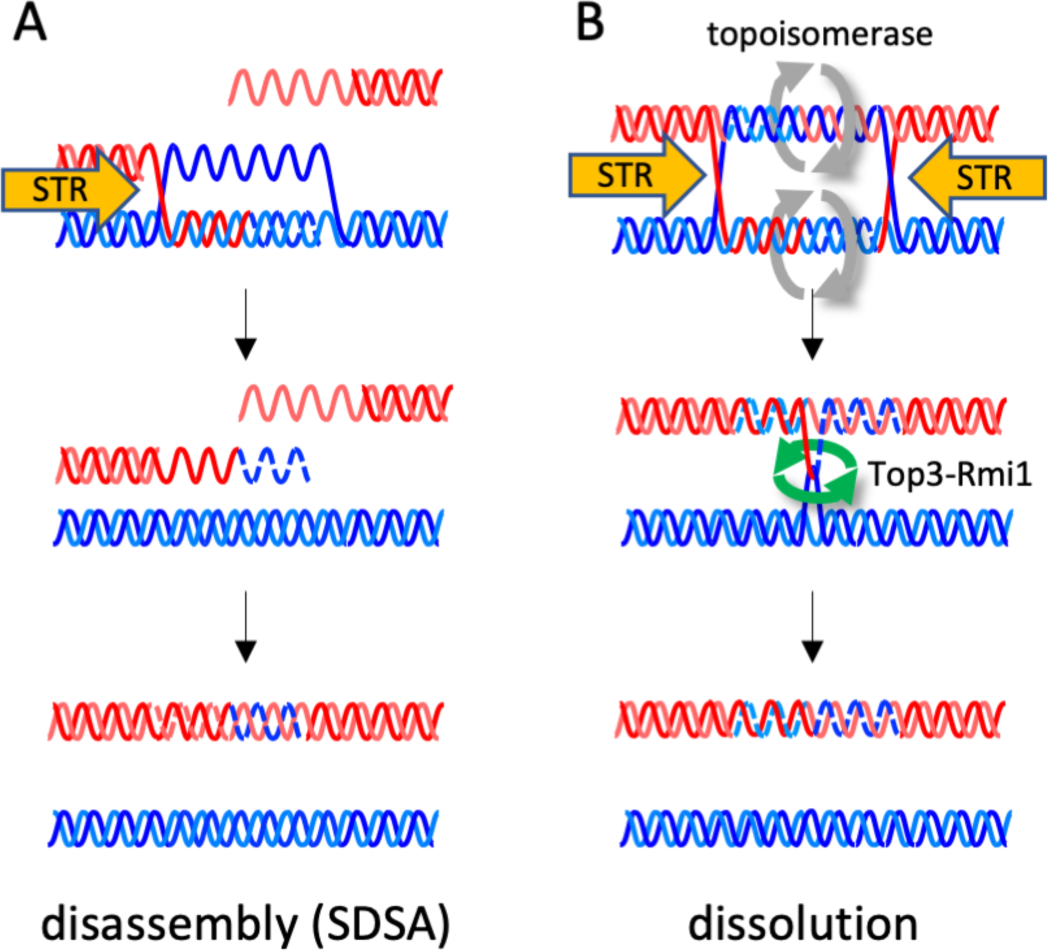
Two possible Sgs1-Top3-Rmi1 anti-crossover activities. (A) D-loop disassembly and synthesis-dependent strand annealing (SDSA). Following strand invasion and 3’ end-primed synthesis (indicated by dashed lines), an unwinding activity (orange arrow, in this case the STR complex) takes apart a D-loop, releasing the invading break end. Annealing with the other break end, followed by gap-filling synthesis, produces a noncrossover recombinant. (B) Dissolution. Unwinding activities drive convergent Holiday junction migration, facilitated by relief of overwinding (gray arrows), to produce two linear DNA molecules linked by at least one hemicatenane. Single-strand passage by Top3-Rmi1 (green arrows) resolves hemicatenanes and produces a noncrossover recombinant.

Consistent with these *in vitro* activities, *sgs1, top3*, and *rmi1* mutants (hereafter referred to collectively as *str* mutants) display elevated levels of mitotic crossing-over and DNA damage sensitivity. *str* mutants are also synthetic lethal with mutants lacking either Mus81-Mms4 or Slx1-Slx4, nucleases that resolve dHJ-JMs (Wallis *et al*. 1989; Mullen *et al*. 2001; Shor *et al*. 2002; Ira *et al*. 2003; Mullen *et al*. 2005; Ehmsen and Heyer 2008; Wyatt and West 2014). This synthetic lethality is suppressed by reducing or eliminating homologous recombination (Fabre *et al*. 2002), consistent with the suggestion that *str* mutants accumulate recombination intermediates that are toxic if unresolved. When exposed to DNA damaging agents, *str* mutants accumulate JMs to levels that are greater than in wild type (Mankouri *et al*. 2011; Ashton *et al*. 2011), again consistent with a role for STR in limiting the accumulation of dHJ-JMs. While these mutant phenotypes are caused by loss of any STR component, it is clear that the Top3-Rmi1 heterodimer has important activities independent of Sgs1. Cells lacking Top3 or Rmi1, but not Sgs1, display slow growth, accumulate G2/M phase cells, and show persistent DNA damage signaling, consistent with low-level induction of the DNA damage response (Wallis *et al*. 1989; Gangloff *et al*. 1994; Chakraverty *et al*. 2001; Chang *et al*. 2005; Mullen *et al*. 2005). Loss of Sgs1 activity or of homologous recombination suppresses these phenotypes (Gangloff *et al*. 1994; Oakley *et al*. 2002; Shor *et al*. 2005; Chang *et al*. 2005), suggesting that Top3-Rmi1 limits the accumulation of toxic recombination intermediates formed by Sgs1. While these findings point to an important role for STR in modulating homologous recombination, they do not distinguish between an early role in D-loop disassembly and a late role in dHJ-JM dissolution.

Support for a role for STR in D-loop disassembly has come from studies of meiotic and mitotic recombination. In budding yeast, most meiotic NCOs are thought to be formed by SDSA, without a stable dHJ-JM intermediate (Allers and Lichten 2001), while most meiotic COs derive from dHJ-JMs that are stabilized by an ensemble of meiosis-specific proteins called the ZMM proteins (Börner *et al*. 2004; Lynn *et al*. 2007; Pyatnitskaya *et al*. 2019) and are resolved as COs by the MutL*γ* (Mlh1, Mlh3, Exo1) complex (Argueso *et al*. 2004; Zakharyevich *et al*. 2010; 2012). Consistent with NCO formation by STR-mediated D-loop disassembly, *str* mutants no longer form meiotic NCOs by SDSA, and instead use a third pathway that involves ZMM-independent JM formation and resolution, as both NCOs and COs, by structure selective nucleases (SSNs; Mus81-Mms4, Yen1, and Slx1-Slx4) that also resolve JMs during the mitotic cell cycle (Oh *et al*. 2007; Matos *et al*. 2011; De Muyt *et al*. 2012; Kaur *et al*. 2015; Tang *et al*. 2015). Thus, in addition to being needed for JM-independent NCO formation during meiosis, the STR complex also determines whether meiotic JMs form in a ZMM-dependent or independent manner. Evidence that STR activity limits the formation of strand-invasion intermediates in mitotic cells has been provided by studies that used a proximity ligation assay to detect early associations between recombining chromosomes during DSB repair (Piazza *et al*. 2019); this signal increased about 2-fold both in *sgs1Δ* mutants and in strains overexpressing a catalysis-dead *top3* mutant protein. Because this study used a repair substrate with homology only to one side of the DSB, it could not directly address the role of the STR complex in modulating dHJ-JM formation.

The studies described above also revealed Sgs1-independent function for Top3-Rmi1 during meiosis. In *top3* and *rmi1* mutants, but not in *sgs1*, a substantial fraction of JMs remain unresolved and impair meiotic chromosome segregation. This indicates that Top3-Rmi1 prevents the accumulation of JMs where the two parental DNA molecules are linked by structures, such as hemi-catenanes (Giannattasio *et al*. 2014), that are not resolved by SSNs (Kaur *et al*. 2015; Tang *et al*. 2015). Interestingly, unlike what is observed in mitotic cells, *sgs1* mutation did not suppress the meiotic JM-resolution defect of *top3* and *rmi1* mutants, indicating that Sgs1 is not responsible for forming these unresolvable intermediates.

I*n vivo* data showing that STR/BTR can resolve mature dHJ-JMs by dissolution has been even more limited, and has come from studies using yeast *ndt80* mutants, which arrest in meiosis I prophase with unresolved JMs. Tang *et al*. (2015) combined conditional-depletion allele of *RMI1* with inducible expression of *NTD80* to study JM resolution under conditions of Rmi1 depletion. Rmi1-depleted cells displayed incomplete JM resolution and were unable to complete meiotic chromosome segregation, consistent with a role for Top3-Rmi1 in the resolution of at least some of the JMs that form during normal meiosis. However, because of ongoing JM formation during the initial stages of Rmi1-depletion, this study could not exclude the possibility that JMs with altered structures, formed under conditions of reduced STR activity, were responsible for the observed resolution and chromosome segregation failures. In a second study, Dayani *et al*. (2011) examined meiotic JM resolution during return-to-growth (RTG). In RTG, cells undergoing meiosis are shifted to rich growth medium, whereupon they exit meiosis and return to the mitotic cell cycle, during which time meiotic JMs are resolved under conditions similar to the G2 phase of the mitotic cell cycle (reviewed in Simchen 2009). JM resolution during RTG was delayed, relative to wild type, in substrate recognition-defective *sgs1-ΔC795* mutants (Schiller *et al*. 2014), consistent with STR promoting early JM resolution by dissolution. However, this study could not exclude the possibility that JMs with altered structures form during meiosis in *sgs1-ΔC795* mutants, and that this structural difference, rather than the absence of active Sgs1, was responsible for the observed delay in resolution.

To further test STR complex-mediated dHJ-JM dissolution *in vivo*, we used an experimental approach that combines RTG with conditional depletion of Sgs1 and/or Rmi1, so that JMs accumulated during meiosis in the presence of normal STR function could then be resolved during RTG in either the presence or absence of active STR. Our findings support a role for STR-mediated JM resolution by dissolution during the mitotic cell cycle, and provide further evidence for an Sgs1-independent Top3-Rmi1 function during JM resolution. In addition, we provide evidence that the DNA damage response prevents cell cycle progression when unresolved recombination intermediates are present.

## Materials and Methods

### Strains

Yeast strains (Table 1) were derived from the haploid parents of MJL2984 (Jessop *et al*. 2005) by genetic crosses or transformation. Transformants were confirmed by PCR and/or Southern blot analysis. All protein fusions were confirmed by sequencing PCR products amplified from the genome.

**Table 1.** Strains used.

| Name | Genotype |
| --- | --- |
| MJL3807 | *URA3::PTEF1-OsTIR1/ URA3::PTEF1-OsTIR1 SGS1-3xHA-IAA17-hygMX/SGS1-3xHA-IAA17-hygMX* |
| MJL3847 | *URA3::PZEO1-OsTIR1/ URA3::PZEO1-OsTIR1 RMI1-AID\*-9xMYC-hphNT1/ RMI1-AID\*-9xMYC-hphNT1* |
| MJL3863 | *URA3::PZEO1-OsTIR1/ URA3::PZEO1-OsTIR1 SGS1-3xHA-IAA17-hygMX/SGS1-3xHA-IAA17-hygMX RMI1-AID\*-9xMYC-hphNT1/ RMI1-AID\*-9xMYC-hphNT1* |
| MJL3899 | *URA3::PZEO1-OsTIR1/ URA3::PZEO1-OsTIR1 RMI1-AID\*-9xMYC-hphNT1/ RMI1-AID\*-9xMYC-hphNT1 PDS1-18xMYC-LEU2/ PDS1-18xMYC-LEU2* |
| S5338xS5344<sup>1</sup> | *URA3::PZEO1-OsTIR1/ ura3 RMI1-AID\*-9xMYC-hphNT1/ RMI1-AID\*-9xMYC-hphNT1 PDS1-18xMYC-LEU2/ PDS1-18xMYC-LEU2 mad1Δ::natMX/mad1Δ::natMX* |
| S5342xS5348<sup>1</sup> | *URA3::PZEO1-OsTIR1/ ura3 RMI1-AID\*-9xMYC-hphNT1/ RMI1-AID\*-9xMYC-hphNT1 PDS1-18xMYC-LEU2/ PDS1-18xMYC-LEU2 rad9Δ::natMX/rad9Δ::natMX* |

<sup>1</sup> *mad1Δ* and *rad9Δ* strains were maintained as haploid stocks, and fresh diploids were isolated before each experiment.

### Return to growth

Induction of meiosis, return to growth, and protein depletion were performed as described (Dayani *et al*. 2011; Kaur *et al*. 2018). Briefly, meiosis was induced in 400 mL liquid cultures at 30°C. After 6 h, cells were harvested by centrifugation, washed with water, resuspended in the same volume of growth medium (YPAD) prewarmed to 30°C, divided equally between two 2 L baffled Erlenmeyer flasks and aerated vigorously (350 rpm) at 30°C. Auxin (indole acetic acid, 0.5 M stock in DMSO) was added to one culture to a final concentration of 2 mM, and the same volume of DMSO was added to the other These additions were repeated every subsequent hour. Samples for DNA, protein, and cytological analysis were collected at indicated time points.

### DNA extraction and analysis

DNA isolation and recombination intermediate and product detection were performed as described (Allers and Lichten 2000; 2001; Oh *et al*. 2009). As illustrated in Figure S1, recombination intermediates were scored on blots of gels containing *Xmn*I digests, probed with *ARG4* coding sequences (+156 to +1413). Crossover and noncrossover products were scored on blots of gels containing *Eco*RI-*Xho*I digests, probed *HIS4* coding sequences (+539 to +719).

### Protein extraction and western blotting

Protein extracts were made by TCA precipitation (Foiani *et al*. 1994) from 3 mL of cultures. Gel electrophoresis, blotting, and probing were performed as described (Kaur *et al*. 2018). Primary antisera and dilutions were: mouse anti-HA monoclonal (clone 12CA5, Roche; 11583816001), 1/10,000; rabbit anti-MYC (Santa Cruz, sc-789), 1/1000; goat anti-ARP7 (Santa Cruz Biotechnology, sc-8961), 1/1000. Secondary antibodies were alkaline phosphatase conjugates of goat anti-mouse IgG (Sigma, A3562); rabbit anti-goat IgG (Sigma, A4187); and goat anti-rabbit IgG (Sigma, A3687). All were used at 1/10,000 dilution.

### Cytology

Cells were prepared for cytological analysis by immunostaining as described (Xaver *et al*. 2013) with the following modifications. Cells were fixed with three successive incubations (10-15 min, room temperature) in 3.4% formaldehyde, the latter two in 0.1M potassium phosphate, 0.5 mM MgCl_2_, pH 6.4, and then were stored at 4°C. Spheroplasting used 0.5 mg/ml Zymolyase 100T (Nacalai USA #07655) in place of Zymolyase 20T. Slides were immunostained overnight at 4°C or 4h at 30°Cwith a mixture of the two primary antisera diluted in blocking buffer [rat anti-tubulin (Abcam ab6160 1:1250) and rabbit anti-MYC (Santa Cruz, sc-789 1:250)], washed in PBS (3 time, 5 min, room temperature), and were then incubated with secondary antisera [Cy3-conjugated donkey anti-rabbit IgG (Jackson Laboratories, #711-165-152) and FITC-conjugated rabbit anti-rat IgG (Sigma, # F1763), both 1:600 in blocking buffer] for 3 h at 30°C, followed by three 5 min room temperature washes in PBS. Samples to be examined by DAPI-staining only were treated as described (Goyon and Lichten 1993) after formaldehyde fixation and storage as above.

### Estimation of unresolved joint molecules

The number of unresolved JMs in Rmi1-depleted cells were estimated using previous calculations of about 90 interhomolog COs per nucleus (Mancera *et al*. 2008; Chen *et al*. 2008; Martini *et al*. 2011) and an intersister : interhomolog JM ratio of about 1:4 (Goldfarb and Lichten 2010), for a total of about 113 JMs/nucleus (90 interhomolog and 22.5 intersister). Since about 20% of JMs remain unresolved at 4h after return to growth in Rmi1-depleted cells (Figure 2B, below), we calculate that about 18 interhomolog JMs and about 4.5 intersister JMs remain unresolved, per nucleus, at 4h after return to growth. The first nuclear division after return to growth involves sister chromatid segregation (Dayani *et al*. 2011), and we presume random segregation of homolog chromatids. All unresolved intersister JMs and ½ of all unresolved interhomolog JMs are expected to be in a configuration that prevents segregation during mitosis; therefore, there would be a minimum of 13 unresolved JMs per nucleus with the potential to prevent chromosome segregation during RTG. Because cells undergoing RTG are tetraploid (Dayani *et al*. 2011), this corresponds to about 40% of all chromatids segregating from each other.

**Figure 2.**
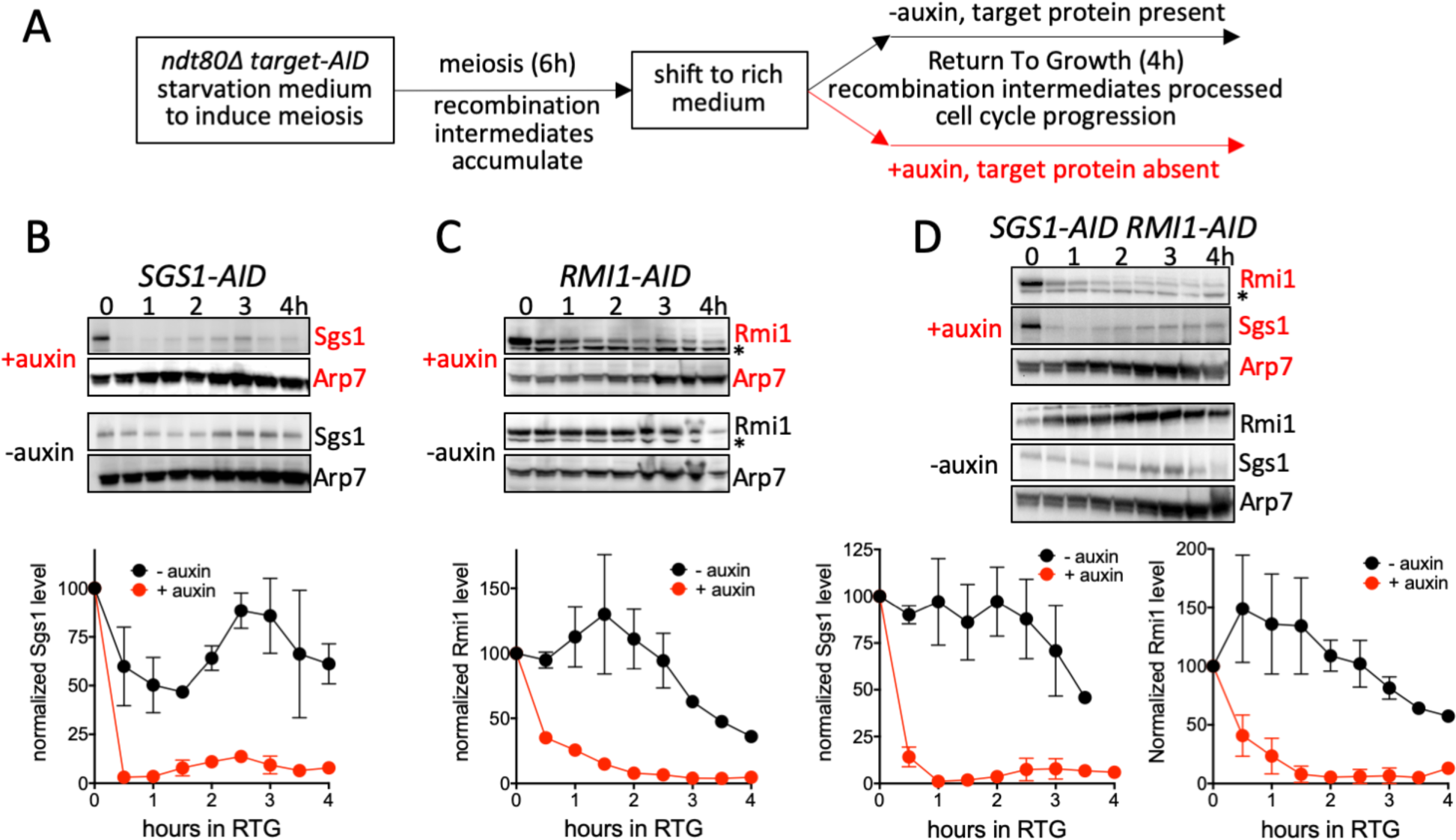
Use of auxin-inducible degrons during return to growth (RTG). (A) Experimental plan. Diploid budding yeast *ndt80Δ* mutants, containing an auxin-inducible degron (AID) fused to the protein of interest and constitutively expressing the rice OsTIR1 auxin response F-box protein, are induced to undergo meiosis and accumulate meiotic recombination intermediates for 6 h. Cells are then shifted to rich medium, at which point they re-enter the mitotic cell cycle, during which cell cycle landmarks, recombination intermediates, and products are monitored. If auxin is present, the protein of interest is degraded; if auxin is absent, the protein of interest remains. (B) Auxin induced degradation of Sgs1-AID (MJL3807). Top: representative Western blot sections, probed with anti-HA to detect Sgs1 and with anti-Arp7 as a loading control. Times are hours after shift to rich medium. Bottom: normalized Sgs1 levels (Sgs1/Arp7, with 0 h set to 100); red—auxin present; black—vehicle only. (C) Auxin-induced degradation of Rmi1 (MJL3847), details as in panel (B), except that anti-Myc was used to detect Rmi1. The lower band in the Rmi1 panels (*) is a cross-reacting protein. (D) Auxin-induced degradation of Sgs1 and Rmi1 in a doubly-tagged strain (MJL3863), details as in panels (B) and (C). Values are the mean of two independent experiments; error bars indicate range.

### Data availability

All experimental materials not supplied commercially will be supplied upon request. Authors affirm that all data necessary to confirm the conclusions of this article are present within the article, figures and tables. Numerical values underlying graphs in all figures are provided in File S1.

## Results

### Targeted degradation of Sgs1 and Rmi1 during return to growth

To study STR function during RTG, we used auxin-mediated protein degradation (Nishimura *et al*. 2009; Kaur *et al*. 2018) to deplete Sgs1 and/or Rmi1 in a controlled manner (Figure 2A). Strains contained Sgs1 and/or Rmi1 fused to an auxin-inducible degron (AID) and OsTIR1, a rice-derived, auxin-specific F-box protein expressed from a strong constitutive promoter (see Table 1). Similar strains containing a Top3-AID fusion did not display consistent Top3 depletion and therefore were not further studied (H. Kaur, unpublished). Strains also contained a deletion of *NDT80*. Ndt80 drives meiotic expression of the Cdc5 polo-like kinase (Chu and Herskowitz 1998), which activates JM resolution in both meiotic and mitotic cells (Clyne *et al*. 2003; Sourirajan and Lichten 2008; Matos *et al*. 2011; Blanco *et al*. 2014).

In experiments performed here, cells underwent meiosis for 6 hours and accumulated JMs in the presence of normal STR complex function. RTG was then initiated by shifting cells from sporulation medium to rich growth medium. Under these conditions, cells rapidly reduce meiotic transcripts, disassemble the synaptonemal complex, repair remaining DSBs, and resume the mitotic cell cycle, including bud emergence and a mitotic cell division (segregating sister chromatids) without an intervening S phase (Zenvirth *et al*. 1997; Friedlander *et al*. 2006; Dayani *et al*. 2011). To examine JM processing and resolution in the absence or presence of Sgs1 and/or Rmi1, either auxin or vehicle were added at the time of shift to growth medium and every hour afterward (Figure 2A). In the presence of auxin, Sgs1-AID levels reduced to background by 1 h after RTG initiation and auxin addition (Figure 2B, D). Rmi1-AID depletion was less rapid, reaching ∼75% of initial levels after 1 h, and background levels at 2 h (Figure 2C, D). This corresponds to the time when buds first emerge and is more that 30 min before the time that the nuclear division is first visible (see Figure 4, below).

**Figure 3.**
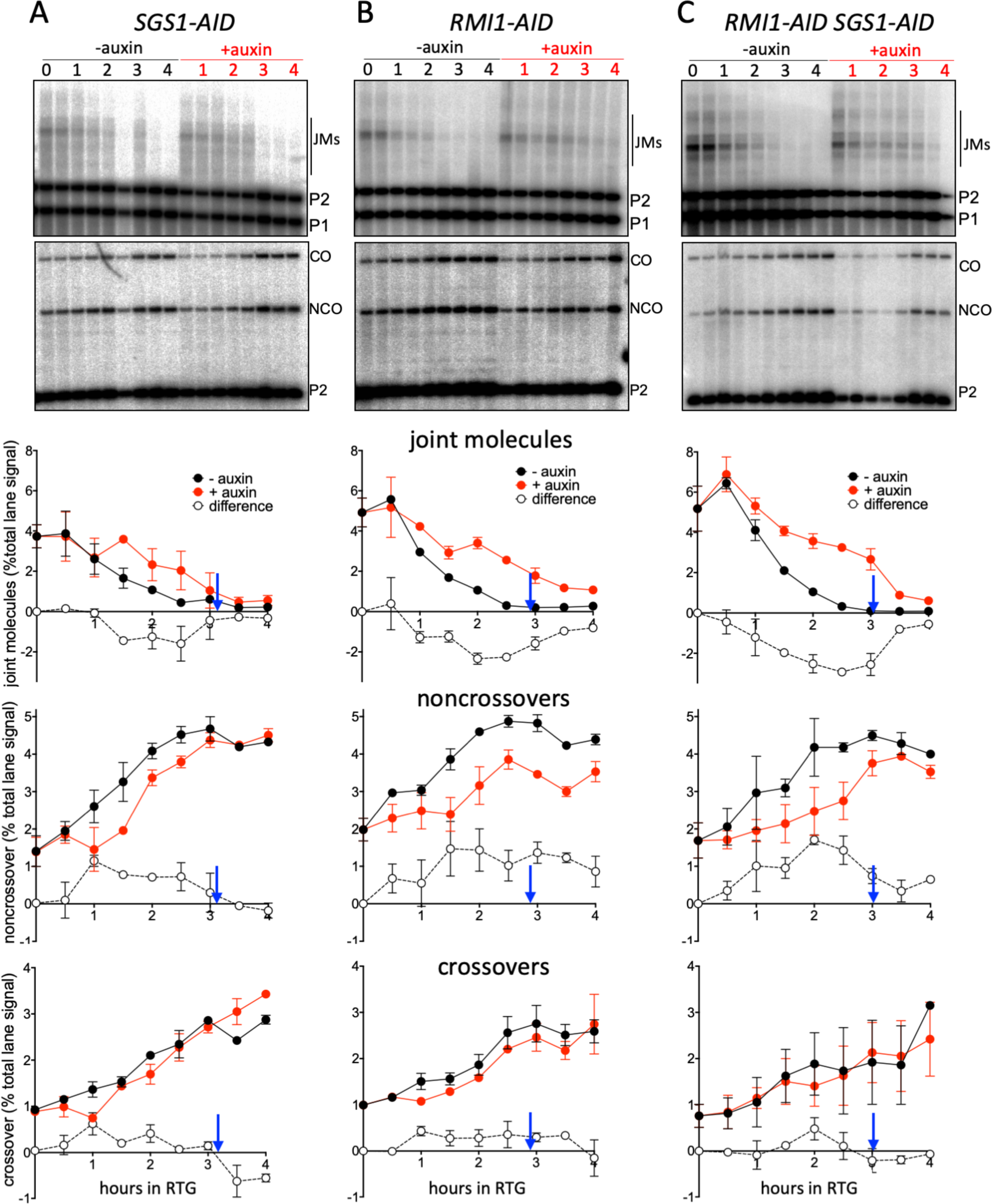
Recombination intermediate resolution and recombination product formation during RTG. DNA was extracted at the indicated times and displayed on Southern blots, using restriction enzymes and probes to detect recombination intermediates (joint molecules, JMs) or crossover (CO) and noncrossover (NCO) recombinants. For details, see Figure S1 and Materials and Methods. (A) *SGS1-AID* (MJL3807). Top two panels: representative Southern blots with *Xmn*I and *Eco*RI/*Xho*I digests, probed to detect joint molecules and recombination products, respectively. Bottom three panels: quantification of JMs, NCOs, and COs, expressed as percent of total lane signal. Red—auxin added; black—vehicle only; open circles—difference between levels when Sgs1 is present (-auxin) and depleted (+auxin). Blue arrows indicate when 50% of control cultures (-auxin) had initiated mitosis (see Figure 4, below). (B) *RMI1-AID* (MJL3847) Details as in panel (A). (C) *SGS1-AID RMI1-AID* (MJL3863). Details as in panel (A). Values are the mean of two independent experiments; error bars indicate range.

**Figure 4.**
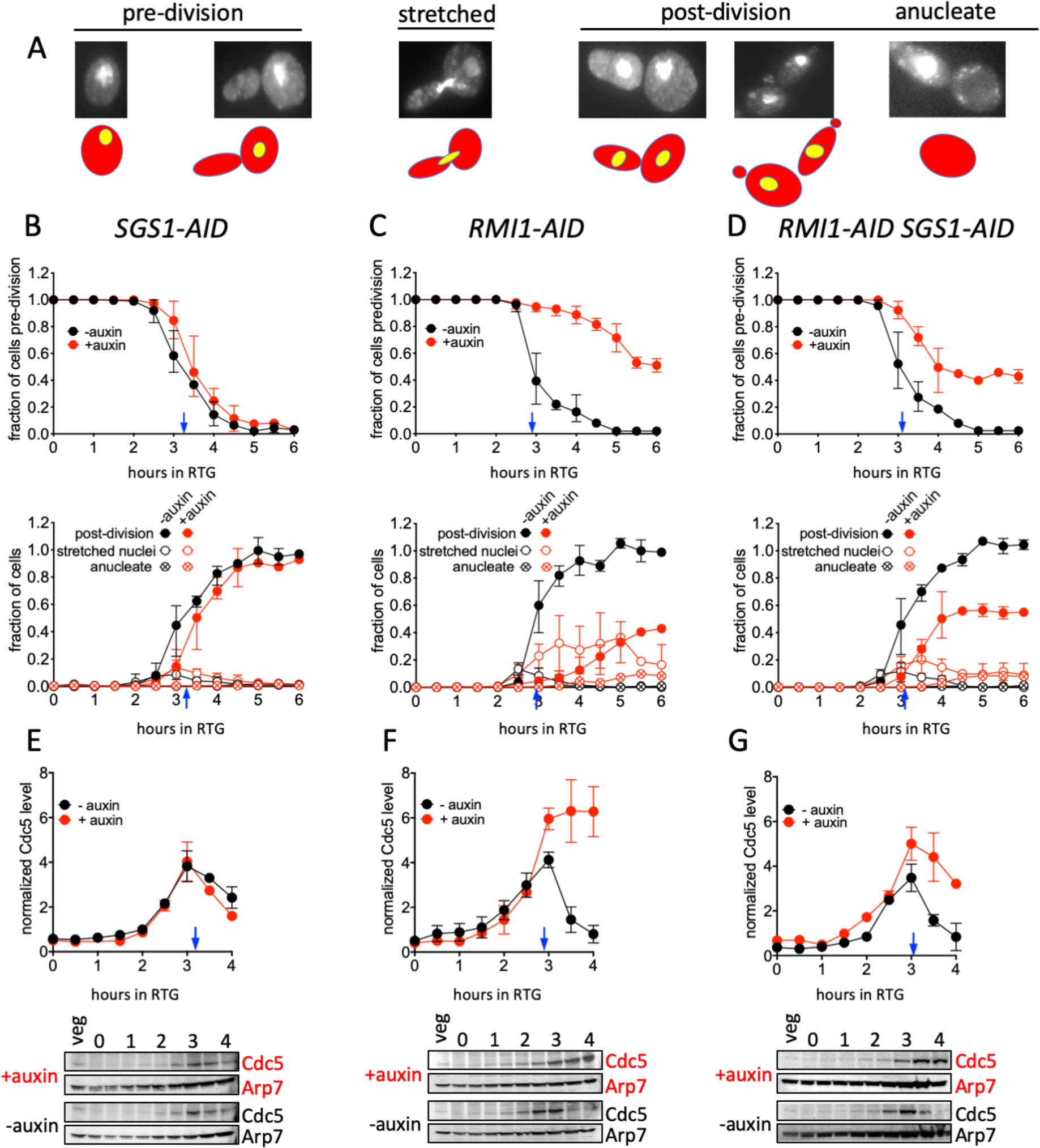
Rmi1-depletion impairs cell cycle progression during RTG. (A) Illustration of cell cycle stages, scored using fixed, DAPI-stained cells. Note that the elongated shape of the first bud to emerge during RTG allows distinction between original mother cells and daughter cells (Dayani *et al*. 2011). “Pre-division”—unbudded cells and cells with a bud and a single nucleus in either the mother or daughter; “stretched”—cells with a single nucleus stretched between mother and daughter; “post-division”— elongated, nucleated cells, with or without a bud; “anucleate”—cells with no nuclear DNA staining but with visible mitochondrial staining. Since the first division after RTG produces one elongated and one round cell, the number of elongated cells can be used to infer the number of round cells produced by this division. Panels (B), (C) and (D)—upper panel, fraction of predivision cells; lower panel cells completing mitosis (“post-division”, solid circles) or in the midst of mitosis (“stretched”, hollow circles) for *SGS1-AID* (MJL3807), *RMI1-AID* (MJL3847) and *SGS1-AID RMI1-AID* (MJL3863), respectively, in control (black) or auxin-mediated depletion (red) conditions. Values from 0 to 4h are the mean of three independent experiments; those from 4.5 to 6h are the mean of two of these three experiments. Error bars indicate range. Panels (E), (F) and (G)—Cdc5 protein levels during RTG in *SGS1-AID* (MJL3807), *RMI1-AID* (MJL3847) and *SGS1-AID RMI1-AID* (MJL3863), respectively. Bottom panels—representative Western blot sections probed for Cdc5 or for Arp7 as a loading control; a sample from an exponentially-growing culture (“veg”) is included to allow blot-to-blot normalization. Top panels— normalized Cdc5 levels, calculated as the Cdc5/Arp7 ratio for experimental time points divided by the Cdc5/Arp7 ratio of the “veg” control. Blue arrows indicate when 50% of control (-auxin) cultures had initiated mitosis. Values are the mean of two independent experiments; error bars indicate range.

### Rmi1 and Sgs1 are needed for timely JM resolution during RTG

To monitor JM processing and resolution during RTG, we used a well-characterized recombination-reporter construct integrated on the left arm of chromosome *III* in which JMs, COs and NCOs can be quantitatively scored on Southern blots (Jessop *et al*. 2005; Figure S1). Depletion of Sgs1 during RTG delayed JM disappearance and NCO formation by 1 h relative to undepleted controls (Figure 3A), confirming previous conclusions that Sgs1 is needed for timely JM resolution during RTG (Dayani *et al*. 2011). Despite this delay, the majority of JMs had disappeared by 3.5-4 h after RTG (12 ± 4% JMs remaining in Sgs1 depleted cells versus 5 ± 2% in undepleted controls, average of 3.5 and 4h ± SD), when most cells had initiated mitosis (Figure 4B), and equivalent final NCO levels were achieved in both conditions. In contrast, depletion of Rmi1 during RTG both delayed and reduced JM resolution and NCO formation. Considerably more JMs remained in Rmi1-depleted cells (21 ± 4% versus 4 ± 2% in undepleted controls) than in Sgs1-depleted cells (p = 0.014, Welch’s t-test). NCOs were similarly reduced, by 25 ± 8% relative to undepleted controls (Figure 3B). These findings indicate that, when Sgs1 is present, Rmi1 is important for timely JM processing and NCO formation.

Chronic loss of Top3 or Rmi1 results in a slow-growth phenotype that is suppressed by *sgs1* loss-of-function mutants (Gangloff *et al*. 1994; Chang *et al*. 2005). To see if the JM resolution and NCO formation defects observed upon Rmi1 depletion are similarly suppressed, we performed RTG experiments in which both Rmi1 and Sgs1 were auxin-depleted (Figure 3C). Sgs1 co-depletion partially suppressed Rmi1 depletion phenotypes: while the fraction of JMs unresolved was indistinguishable from those in Sgs1 depletion alone (12 ± 3% versus 12 ± 4%), JMs disappearance was slower, with a partial defect in NCO formation. Final NCO levels in Sgs1/Rmi1 co-depleted strains (13 ± 7% reduced relative to undepleted controls) were intermediate between Sgs1-depletion alone (0 ± 9%) and Rmi1-depletion alone (25 ± 8%). Possible reasons for this intermediate phenotype will be discussed below.

### Rmi1 depletion causes DNA segregation and cell cycle-progression defects during RTG

Unresolved JMs formed during meiosis impede nuclear division without affecting other steps of meiotic progression, such as spindle assembly/disassembly and spore wall formation (Jessop and Lichten 2008; Oh *et al*. 2008; De Muyt *et al*. 2012; Kaur *et al*. 2015; Tang *et al*. 2015). To see if similar defects occur during RTG of Rmi1-depleted cells, we monitored nuclear divisions (Figure 4) in the same cultures used to analyze JMs and recombinant products, taking advantage of the fact that the first cell cycle after RTG, unlike subsequent cell cycles, produces elongated buds and daughter cells (Dayani *et al*. 2011). Cells were scored as pre-division (either round unbudded or round mother with an elongated bud and a single nucleus in the mother), as post-division (elongated cells containing a single nucleus) or as in metaphase or anaphase (an undivided nucleus either in the bud neck or stretched between a round mother and elongated daughter; hereafter referred to as “stretched”). In control cultures, cells undergoing mitosis were first seen at 2.5 h after RTG, and virtually all cells had completed mitosis by 5 h, with only a small fraction in metaphase/anaphase at any given time (Figure 4B-D). Cultures depleted for Sgs1 alone also initiated and completed mitosis in a timely manner, albeit with a slight delay (Figure 4B). In contrast, most Rmi1-depleted cells failed to complete mitosis (Figure 4C). A substantial fraction of Rmi1-depleted cells contained “stretched” nuclei at 5h after RTG, a time when mitosis was complete in control cultures. Upon further incubation, this fraction declined, and post-division cells lacking a nucleus (anucleate cells) appeared at low levels. Cultures co-depleted for Sgs1 and Rmi1 displayed an intermediate phenotype (Figure 4D). About half appeared to execute mitosis with timing similar to controls, while the rest failed to divide. As in Rmi1-depleted cultures, a substantial fraction of cells that failed to divide contained “stretched” nuclei, and anucleate cells appeared at low levels upon continued outgrowth. This mixed phenotype parallels the partial defects seen in molecular analyses (Figure 3C, above). Together, the cytological and molecular phenotypes of Sgs1/Rmi1 co-depleted cells suggests that these cultures are heterogeneous, with Rmi1 depletion-induced defects being suppressed in only about half of cells.

### Progression defects in Rmi1-depleted cells are due to a cell cycle arrest

We considered two possible reasons for the failure of Rmi1-depleted cells to complete the first mitosis after RTG. The first is that a mechanical barrier, created by unresolved JMs, prevents nuclear division. If this were the case, cells would be expected to progress through mitosis but might not divide chromosomes between mother and daughter cells. Alternatively, it is possible that unresolved JMs or other DNA structures, formed in the absence of Rmi1, are recognized by a checkpoint system that prevents cell cycle progression.

As an initial test, we monitored levels of the Cdc5 polo-like kinase, which is required for full SSN activity during late G2 and mitosis (Matos *et al*. 2011; 2013). Cdc5 is produced during G2/M (Cho *et al*. 1998) and is degraded upon exit from mitosis (Visintin *et al*. 2008). Cdc5 was first detectable at 1.5-2 h after initiation of RTG. In control cultures and Sgs1-depleted cultures, Cdc5 accumulated until about 3 h, when about half of the cells had initiated mitosis. Cdc5 levels then declined, consistent with these cells exiting mitosis and initiating a second cell cycle (Figure 4E-G). In contrast, in Rmi1-depleted cultures, Cdc5 accumulated to greater levels and never declined (Figure 4F), consistent with a block before exit from mitosis. In cultures that were doubly-depleted for Sgs1 and Rmi1, Cdc5 also accumulated and declined, but to levels that greater than those seen in control cultures (Figure 4G), consistent with the previous inference of culture heterogeneity. Because of the more profound effects seen Rmi1-depleted cultures, and because of the complications inherent in the analysis of heterogeneous cultures, we focused our further efforts on characterizing the arrest seen during RTG in conditions of Rmi1-depletion alone.

To further characterize this arrest, we monitored spindle morphology and Pds1 protein levels (Figure 5). Pds1, the budding yeast securin, accumulates in nuclei during G2 and metaphase, and is degraded at the metaphase-anaphase transition (Cohen-Fix *et al*. 1996). Control cultures displayed all the hallmarks of cells progressing unimpeded through mitosis, including bud formation, a transition from G2/metaphase (cells with bipolar spindles and intranuclear Pds1) to anaphase/post-anaphase (cells with bipolar spindles but lacking intranuclear Pds1), and mother-bud separation (Figure 5B). In contrast, Rmi1-depleted cultures rarely underwent mother-bud separation, and the vast majority of cells contained bipolar spindles and intranuclear Pds1, consistent with a G2/M phase cell cycle arrest (Figure 5C). Taken together, these data indicate that Rmi1 depletion during RTG results in both incomplete JM resolution and cell cycle arrest before the metaphase-anaphase transition.

**Figure 5.**
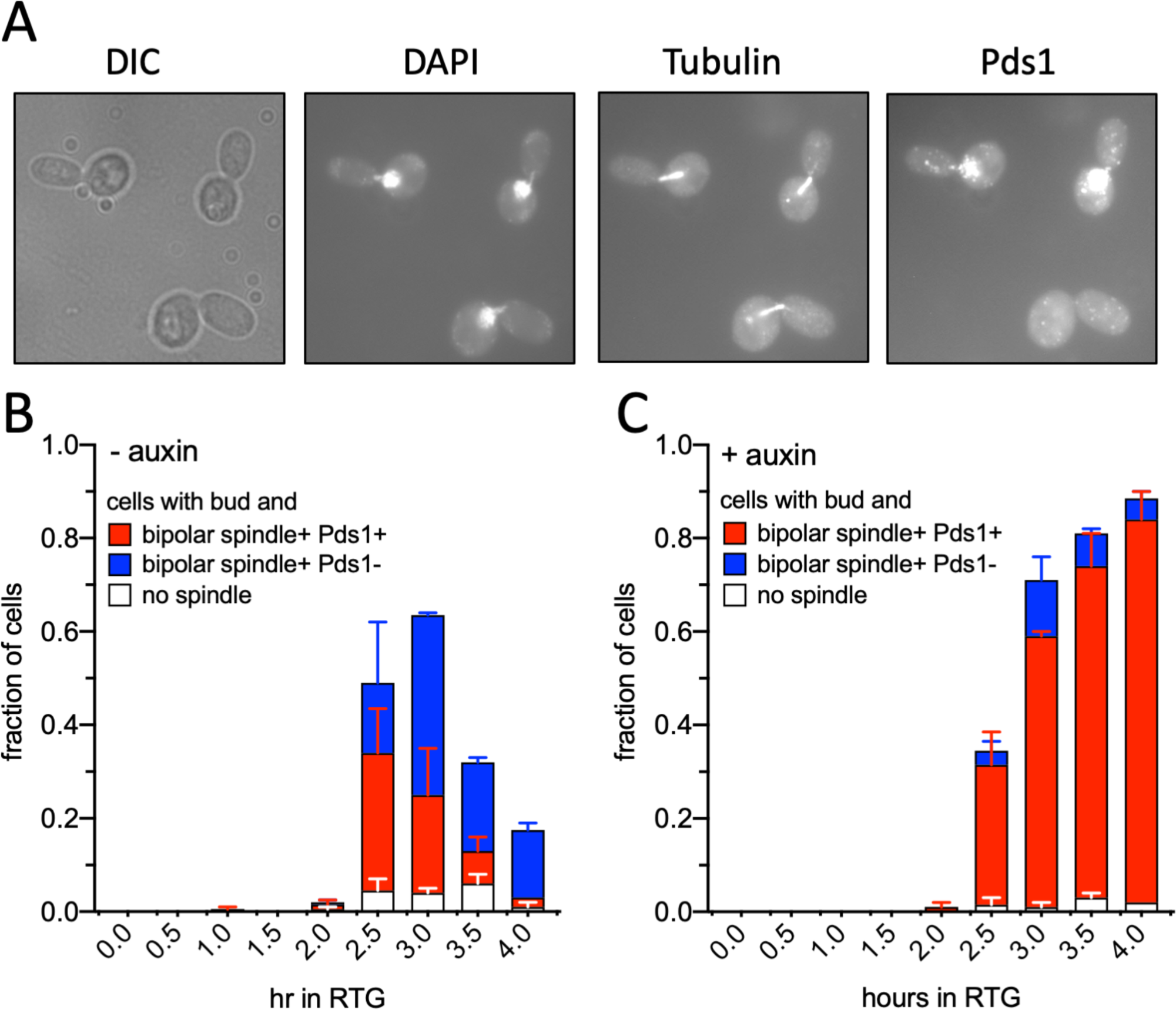
Rmi1-depletion causes a G2/M arrest during RTG. (A) Three Rmi1-depleted mother-daughter cell pairs from an auxin-treated *RMI1-AID* culture (MJL3899) taken 4 h after shift to rich medium containing auxin. From left to right, differential interference contrast image, detection of DNA (DAPI), beta-tubulin, and Pds1-Myc. See Materials and Methods for details. The bottom mother-daughter pair was scored as having undergone the metaphase-anaphase transition, based on the absence of Pds1. (B) Percent of total cells in a control culture with a bud and lacking a bipolar spindle (white) or containing a bipolar spindle and nuclear Pds1 (red) or with a bipolar spindle but lacking nuclear Pds1 (blue). (C) As in panel (B), but in the presence of auxin. Data are from two experiments, error bars denote range.

### Cell cycle arrest during RTG in Rmi1-depleted cells is mediated by the DNA damage response

Two major cell cycle checkpoint systems block Pds1 degradation and cause a G2/M cell cycle arrest: the spindle assembly checkpoint, which detects the presence of kinetochores that are not attached to spindle microtubules (Wells 1996; Cohen-Fix *et al*. 1996); and the DNA damage checkpoint, which detects unrepaired DNA damage (Cohen-Fix and Koshland 1997; Agarwal *et al*. 2003; Harrison and Haber 2006). To determine which system is blocks progression during RTG in the absence of Rmi1, we deleted either *MAD1* or *RAD9*, which are essential for the spindle assembly and DNA damage checkpoints, respectively (Figure 6). When Rmi1 was present, both *mad1Δ* and *rad9Δ* mutants underwent RTG with wild-type efficiency and kinetics, with >90% of cells completing nuclear and cellular division by 4 h after RTG. Rmi1-depleted *mad1Δ* cells displayed arrest phenotypes similar to those seen in *MAD1* Rmi1-depleted cells. Only 9% of cells completed mitosis by 4 h after RTG, and a large fraction of cells (∼40%) contained nuclei with chromosomal DNA stretched between mother and daughter (Figure 6A). In contrast, more than half (57%) of Rmi1-depleted *rad9Δ* mutants completed the first cell division after RTG (Figure 5B). Thus, the DNA damage checkpoint is responsible for the observed cell cycle arrest.

**Figure 6.**
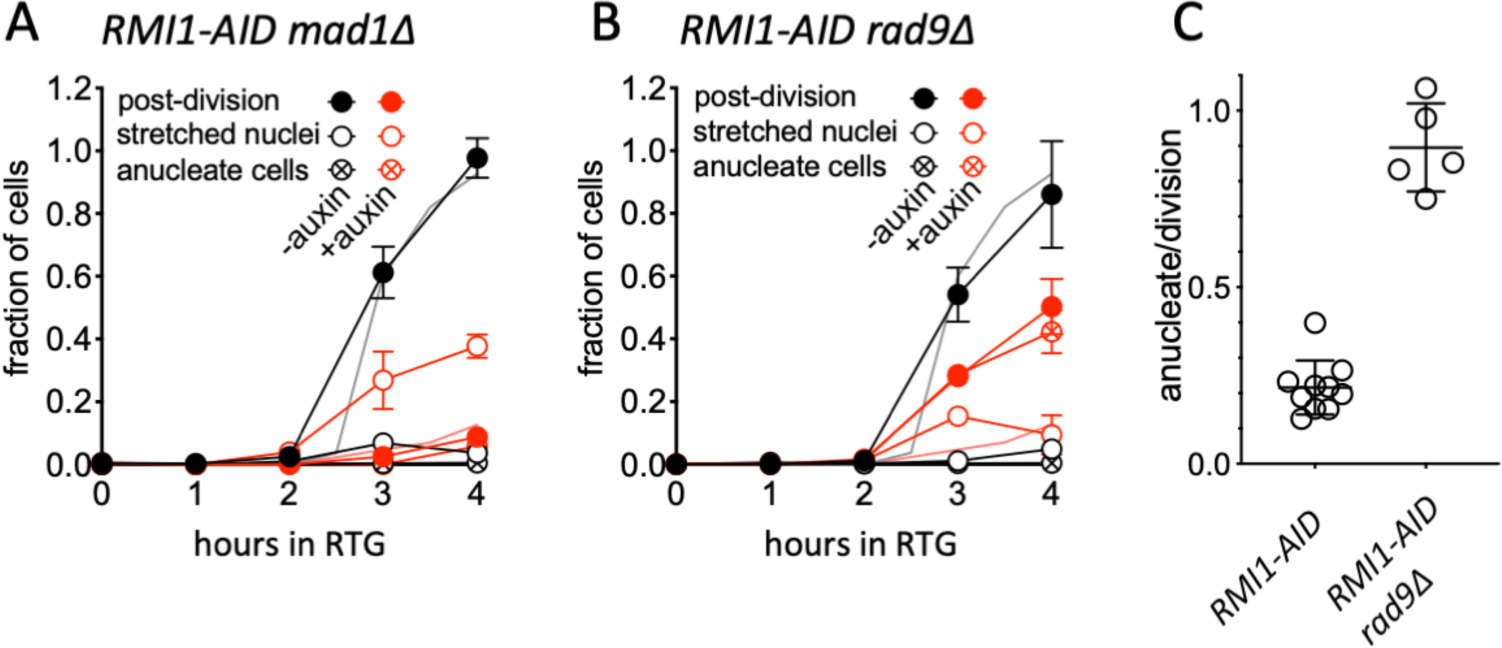
The DNA damage checkpoint is responsible for arresting cell cycle progression during RTG in the absence of Rmi1. (A) Fraction of cells completing cell division (solid circles), at metaphase/anaphase (with chromosomal DNA “stretched” between mother and daughter; hollow circles), or without nuclei (circles with cross) in spindle assembly checkpoint-defective *RMI1-AID mad1Δ* cells (S5338xS5344) during RTG in the absence (black) or presence (red) of auxin. (B) As in panel (A), but with DNA damage checkpoint-defective *RMI1-AID rad9Δ* diploids (S5342xS5348). In both panels A and B, grey and pink lines without symbols are post-division values for corresponding *MAD1 RAD9* diploids, from Figure 4C. (C) Fraction of divisions producing an anucleate cell. Values are from the following time points of two independent experiments: *RMI1-AID*, 4-6h, *RMI1-AID rad9Δ*, 2-4h.

While *rad9Δ* restored cell cycle progression to Rmi1-depleted cells, it did not restore normal chromosome segregation. Instead, at 4h after RTG, about 40% of cells lacked a detectable nucleus (Figure 6B). This corresponds to about 90% of divisions producing an cell (either mother or daughter) without a nucleus (Figure 6C). This stands in contrast to the much lower level of anucleate cells produced in *RAD9* Rmi1-depleted cultures (20% of divisions), where the majority of cells remained arrested. It suggests that unresolved recombination intermediates are present in Rmi1-depleted cells at levels sufficient to mechanically block chromosome segregation when the arrest is bypassed in *rad9Δ* mutants.

## Discussion

### STR-mediated dissolution is an important resolution mechanism during RTG

*In vitro* studies have identified two Sgs1/BLM-Top3-Rmi1 activities, D-loop disassembly (van Brabant *et al*. 2000; Bachrati *et al*. 2006; Fasching *et al*. 2015) and dHJ dissolution (Wu and Hickson 2003; Wu *et al*. 2006; Plank *et al*. 2006), that can limit JM and crossover accumulation. Because most *in vivo* studies have scored either CO end-products or steady-state JM levels, they could not determine if STR/BTR prevents dHJ-JM formation, or if it drives dHJ-JM resolution as NCOs. In the current study, we focused directly on dHJ resolution during RTG under conditions of Sgs1 and/or Rmi1 depletion. Our findings indicate that STR-mediated dissolution is an important mode for dHJ-JM resolution *in vivo*, and that Top3-Rmi1 has important STR-independent functions (see Figure 7).

**Figure 7.**
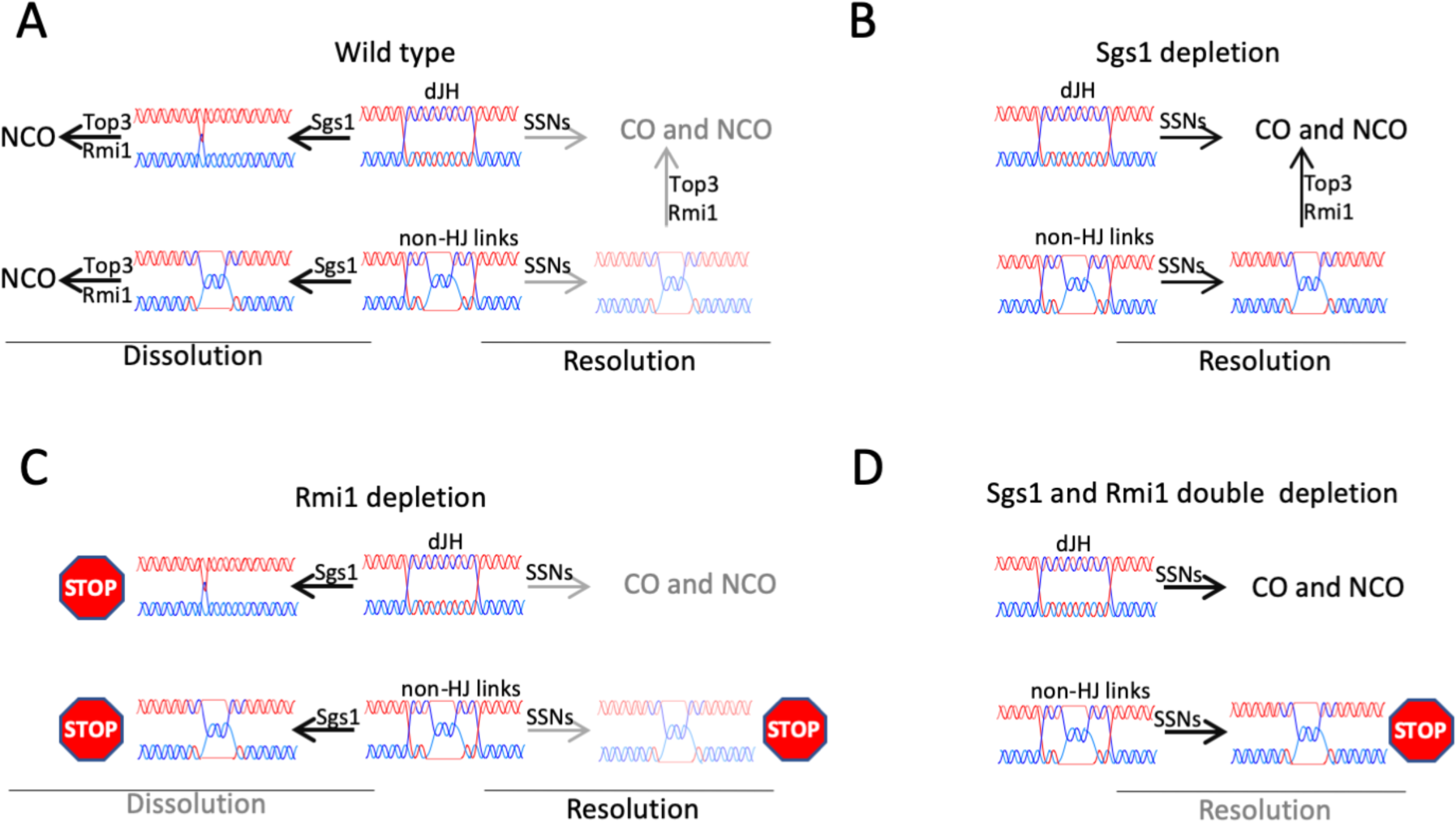
Recombination intermediate resolution during RTG. (A) In cells where the STR is fully functional, most recombination intermediates are resolved by convergent branch migration, shown as involving Sgs1, but which also may involve Top3-Rmi1. The resulting molecules are proposed to contain single-strand interlinks (hemicatenanes), which Top3-Rmi1 resolves to form noncrossovers. A minor fraction of recombination intermediates escape Sgs1 activity and are processed by Holliday junction-resolving nucleases (SSNs). Those that contain only Holliday junctions (top row) are fully resolved, while those containing both Holliday junctions and other strand interlinks (here illustrated as hemicatenanes) require both SSNs and Top3-Rmi1 to produce fully resolved products. (B) In the absence of Sgs1, Holliday junctions in recombination intermediates can be cleaved by SSNs, but intermediates that contain non-HJ interlinks require Top3-Rmi1 to be fully resolved. (C) In the absence of Rmi1, Sgs1-catalyzed convergent branch migration produces molecules with structures that cannot be further resolved, and which induce a Rad9-dependent cell cycle arrest. As in wild-type, some intermediates escape Sgs1 activity and are resolved by SSNs; those containing non-HJ interlinks remain unresolved and, upon stretching, the ssDNA they contain contributes to the cell cycle arrest. (D) In the absence of both Sgs1 and Rmi1, recombination intermediates that contain only Holliday junctions can still be efficiently resolved, while intermediates that also contain non-HJ interlinks will remain unresolved and induce a cell cycle arrest. This figure ignores the possibility that non-STR activities also may contribute to branch migration and decatenation.

dHJ-JMs can be resolved during the mitotic cell cycle either by dissolution or by SSN-mediated HJ cleavage; only the latter can produce COs (Matos and West 2014). We find that, during RTG, Sgs1 or Rmi1 depletion markedly delayed both JM resolution and NCO formation, without changing the timing or levels of COs formed (Figure 3). In addition, even when STR is active, most JM resolution precedes Cdc5 expression and thus SSN activation (Figures 2 and 3). Thus, our findings are consistent with STR-mediated dHJ dissolution being the primary mode of JM resolution during RTG during the mitotic cell cycle.

Still remaining to be answered is question of how JMs are resolved when Sgs1 and/or Rmi1 are deplete. In the absence of STR activity, JMs not resolved by dissolution should be resolved by SSN-mediated cutting in late G2 and mitosis, and thus should produce fewer NCOs and more COs. Our data only partially support this expectation, as NCO formation is delayed when Sgs1 is depleted (Figure 3A, C), but COs do not increase. We consider two possible explanations for this result. First, auxin-mediated Sgs1 depletion may not completely eliminate STR activity and the remaining active fraction might resolve JMs by dissolution before SSNs are activated. Alternatively, Top3-Rmi1 may promote JM dissolution in the absence of Sgs1, as has been reported for D-loop disassembly *in vitro* and *in vivo* (Fasching *et al*. 2015; Piazza *et al*. 2019), either by itself or in combination with other helicases.

### Rmi1 is required for full JM resolution and cell cycle progression during RTG

Previous studies suggest that Top3-Rmi1 has Sgs1-independent functions during the mitotic cell cycle and during meiosis (Wallis *et al*. 1989; Gangloff *et al*. 1994; Chang *et al*. 2005; Mullen *et al*. 2005; Kaur *et al*. 2015; Tang *et al*. 2015), in particular to limit accumulation of JMs that cannot be resolved by standard HJ resolvases. We found that Rmi1 depletion during RTG both delayed and impaired JM resolution and NCO formation (Figure 3B), with unresolved JMs remaining even when Cdc5 levels were high and SSN resolvases should have been fully activated (Figure 4). This is consistent with the suggestion that during the mitotic cell cycle, as in meiosis, Top3-Rmi1 removes inter-molecular DNA connections that cannot be cleaved by HJ resolvases (Kaur *et al*. 2015; Tang *et al*. 2015).

The suppression of *top3* and *rmi1* slow-growth phenotypes by *sgs1* or recombination mutants has led to the suggestion that toxic recombination intermediates are formed by Sgs1 when Top3-Rmi1 is absent (Gangloff *et al*. 1994; Oakley *et al*. 2002; Shor *et al*. 2005; Chang *et al*. 2005). In our study, Sgs1 co-depletion only partially suppressed Rmi1 depletion-associated defects (Figures 3C, 4D). This might have been due to residual Sgs1 activity, possibly present in only some of the cells in the population. However, it is also possible that “toxic” JMs are formed during normal meiosis that require Top3-Rmi1 for their resolution (c.f. Tang *et al*. 2015), even in the complete absence of Sgs1. Persistence of these JMs In Sgs1/Rmi1 co-depleted cells, possibly at levels that vary from cell to cell, might explain the heterogeneous phenotypes of Sgs1/Rmi1 co-depleted cells. Regardless of which explanation is correct, the more penetrant defects observed when Rmi1 alone is depleted support previous suggestions that Sgs1 activity creates JMs that require Top3-Rmi1 for their resolution.

### A DNA damage response-dependent cell cycle arrest during RTG in the absence of Rmi1

Rmi1 depletion during RTG causes a cell cycle arrest at the metaphase-anaphase transition (Figures 4C and 5C) that is bypassed by *rad9Δ*, indicating that it is due to the DNA damage checkpoint (Figure 6B). Remarkably, when this checkpoint is bypassed, almost all of the cells that progress produce an anucleate cell, consistent with unresolved JMs preventing bulk chromosome segregation. We estimate that about 40% of segregating chromatid pairs are linked by unresolved JMs in Rmi1-depleted cells (see Materials and Methods). While this might not be enough to completely block chromosome segregation, the above estimate is based on frequencies of JMs that migrate as discrete species in gels (Figures 3 and S1). The persistent lane background seen in Rmi1-depleted cultures (Figure 3B) may reflect the presence of additional unresolved intermediate structures that might have contributed additional inter-chromatid connections. Further studies will be required to determine the precise nature of these segregation-blocking connections, and of the structures that induce the *RAD9*-dependent DNA-damage checkpoint. Consistent with our finding of a DNA damage checkpoint induced by unresolved JMs during RTG, previous studies have shown that *top3* and *rmi1* mutants display many hallmarks of low-level activation of the DNA damage response (Gangloff *et al*. 1994; Chakraverty *et al*. 2001; Chang *et al*. 2005). Evidence for a DNA damage checkpoint induced by unresolved recombination intermediates is also provided by a report that *sgs1Δ mms4-14A* and *sgs1Δ cdc5-2* mutant strains, which do not activate the Mus81-Mms4 resolvase, also contain an elevated fraction of cells in G2/M (Matos *et al*. 2013).

The DNA damage response is initiated when the Mec1-Ddc2 (ATR-ATRIP) checkpoint kinase interacts with replication protein A-coated single stranded DNA present at unrepaired DNA lesions; Mec1 then acts through intermediary sensors and effectors, including Rad9, to cause cell cycle arrest (reviewed in Nyberg *et al*. 2002). How might unresolved recombination intermediates present during RTG activate the DNA damage response? Little if any break-associated single-strand DNA is expected to be present, since most meiotic DSBs are repaired before the shift to rich medium, especially in *ndt80Δ*-arrested cells, and the few DSBs that remain are rapidly repaired after RTG (Dayani *et al*. 2011).

We suggest that the DNA damage response is induced during RMI-depleted RTG, or in *sgs1Δ* cells unable to activate Mus81-Mms4, when unresolved intermediates that remain undergo stretching by the mitotic spindle, thus exposing single-strand DNA (Figure 7). This in turn raises the question of why similar behavior is not seen during budding yeast meiosis, where meiotic division proceed in the presence of unresolved JMs (Jessop and Lichten 2008; Oh *et al*. 2008; Kaur *et al*. 2015; Tang *et al*. 2015), or during mitosis in mammalian cells, where cells with unresolved links between sister chromatids proceed to anaphase, forming ultrafine DNA bridges (Chan and Hickson 2011; Chan *et al*. 2018). The answer to this question may lie in the different ways that the DNA damage checkpoint functions in mitotically cycling budding yeast, on one hand, and in meiotic yeast and in mammalian cells, on the other. During the budding yeast mitotic cell cycle, chromosomes are always attached to the spindle (Winey and Bloom 2012), and the DNA damage checkpoint blocks the metaphase to anaphase transition (Nyberg *et al*. 2002). Thus, spindle-mediated stretching of unresolved recombination intermediates has the potential to form checkpoint-inducing ssDNA. During budding yeast meiosis, the DNA damage checkpoint blocks expression of the Ndt80 transcription factor that is required for formation of the metaphase I spindle (Winter 2012; Subramanian and Hochwagen 2014; Tsubouchi *et al*. 2018). Once cells activate Ndt80 expression, they are irreversibly committed to undergo meiotic divisions and thus will proceed through meiosis even if unresolved JMs are present (Winter 2012). In a similar vein, the DNA damage response in mammalian cells primarily blocks progression before chromosomes attach to the spindle (Nyberg *et al*. 2002), and multiple mechanisms limit DNA damage response signaling once cells have entered mitosis (Heijink *et al*. 2013). Thus, in both situations, ssDNA would not form at unresolved JMs until it was too late to prevent chromatid separation. Thus, a DNA damage response-mediated cell cycle arrest provoked by unresolved recombination intermediates may be a specific feature of organisms that undergo closed mitosis, and in which chromosomes are always attached to the spindle.

### Concluding remarks

In this work, we have presented data indicating that Sgs1(BLM)-Top3-Rmi1 mediated dissolution is a predominant mechanism for recombination intermediate resolution during the mitotic cell cycle, thus providing *in vivo* confirmation of a mechanism previously proposed by *in vitro* biochemical studies. Our findings also confirm previous suggestions that, in the absence of Top3-Rmi1 decatenase activity, Sgs1 helicase creates entangled structures that cannot be resolved by Holliday junction-cleaving nucleases; similar structures may also be present, albeit at lower levels, in recombination intermediates that form when STR is fully active. Even though all DNA strands in these structures are expected to be intact, our data suggests that their presence activates the DNA damage checkpoint. This unresolved recombination intermediate checkpoint, which perhaps is unique to the budding yeast cell cycle, may be responsible for the observed recombination- and Sgs1-dependent slow growth and G2/M accumulation of *top3* and *rmi1* mutants, and will be fertile ground for future investigation.

## Acknowledgements

This work is dedicated to the memory of Krishnaprasad GN, whose contributions were cut short at the beginning of a promising career. For strain donations and construction, we thank Shangming Tang and Neil Hunter (AID-tagged Rmi1), Takashi Kubota and Anne Donaldson (yeast codon-optimized OsTIR1), Angelika Amon (Pds1-18myc), and Jonathan Staben (Sgs1-3xHA-IAA17). We thank Orna Cohen-Fix for discussions and Jasvinder Ahuja, Matan Cohen, Julia Cooper, and Anura Shodhan for suggestions that improved the manuscript. This work was supported by the Intramural Research Program at the Center for Cancer Research, National Cancer Institute, National Institutes of Health.

---

**Figure S1.**
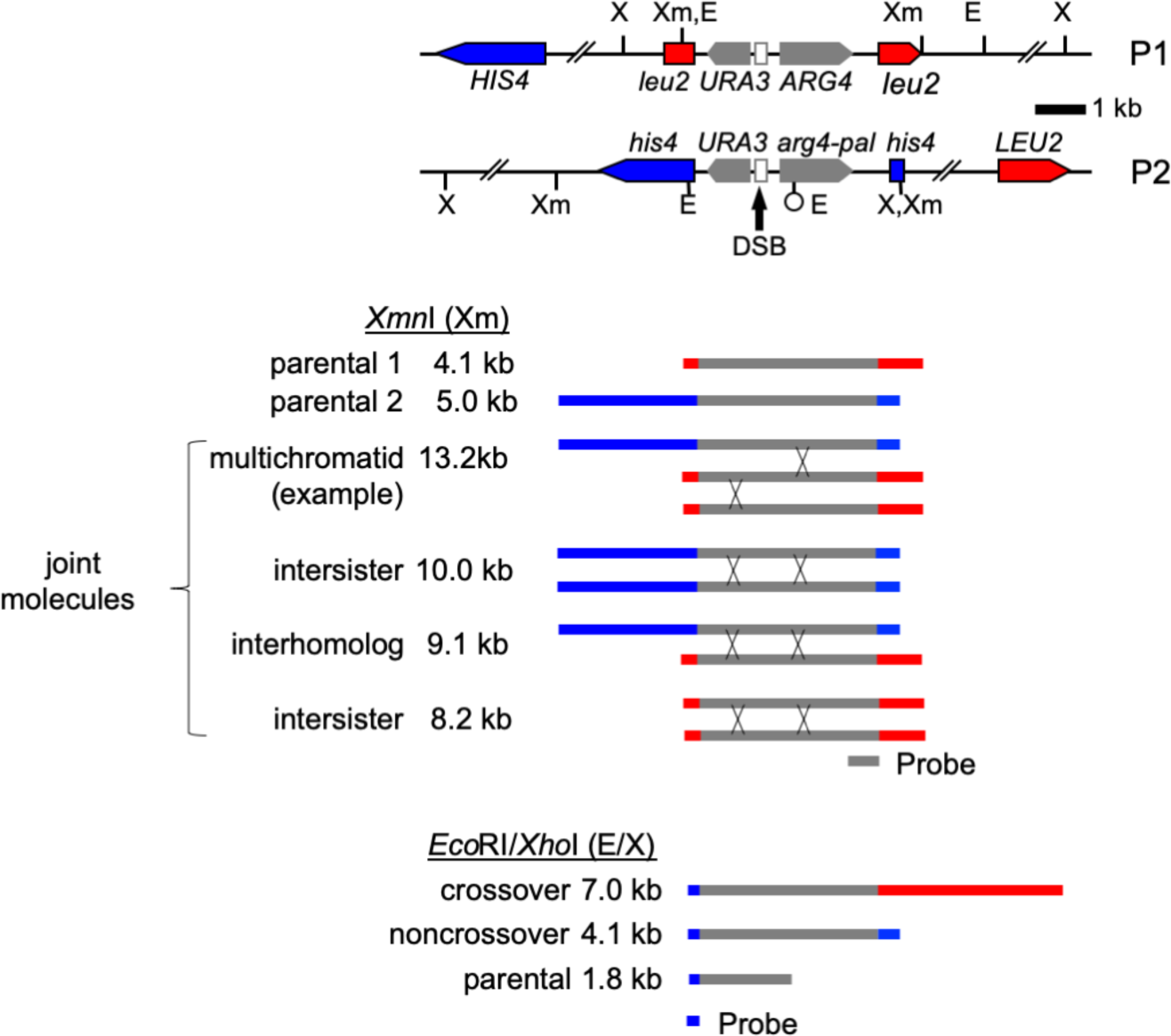
Recombination reporter system used to score recombination intermediates and products (Jessop *et al*. 2005). Diploid strains contain a divergently oriented URA3-ARG4 inserted at the *LEU2* locus on one copy of chromosome *III* (P1), and a similar construct with an *Eco*RI site-marked short palindrome inserted in *ARG4* (*arg4-pal*) at the *HIS4* locus on the other copy of chromosome *III* (P2). *Eco*RI-*Xho*I double digests produce restriction fragments diagnostic of crossovers and of noncrossovers where *arg4-pal* is gene converted to *ARG4*, while recombination intermediates, including interhomolog, intersister, and multichromatid joint molecules, are detected using *Xmn*I digests.

## References

Agarwal R., Tang Z., Yu H., Cohen-Fix O., 2003 Two distinct pathways for inhibiting Pds1 ubiquitination in response to DNA damage. J Biol Chem 278: 45027–45033.

Allers T., Lichten M., 2000 A method for preparing genomic DNA that restrains branch migration of Holliday junctions. Nucleic Acids Res 28: e6.

Allers T., Lichten M., 2001 Differential timing and control of noncrossover and crossover recombination during meiosis. Cell 106: 47–57.

Argueso J. L., Wanat J., Gemici Z., Alani E., 2004 Competing crossover pathways act during meiosis in *Saccharomyces cerevisiae*. Genetics 168: 1805–1816.

Ashton T., Mankouri H., Heidenblut A., Mchugh P., Hickson I., 2011 Pathways for Holliday junction processing during homologous recombination in *Saccharomyces cerevisiae*. Mol Cell Biol 31: 1921–1933.

Bachrati C. Z., Borts R. H., Hickson I. D., 2006 Mobile D-loops are a preferred substrate for the Bloom’s syndrome helicase. Nucleic Acids Res 34: 2269–2279.

Bennett R. J., Sharp J. A., Wang J. C., 1998 Purification and characterization of the Sgs1 DNA helicase activity of *Saccharomyces cerevisiae*. J Biol Chem 273: 9644–9650.

Bernstein K. A., Gangloff S., Rothstein R., 2010 The RecQ DNA helicases in DNA repair. Annu Rev Genet 44: 393–417.

Blanco M. G., Matos J., West S. C., 2014 Dual control of Yen1 nuclease activity and cellular localization by Cdk and Cdc14 prevents genome instability. Mol Cell 54: 94–106.

Börner G. V., Kleckner N., Hunter N., 2004 Crossover/noncrossover differentiation, synaptonemal complex formation, and regulatory surveillance at the leptotene/zygotene transition of meiosis. Cell 117: 29–45.

Cejka P., Cannavo E., Polaczek P., Masuda-Sasa T., Pokharel S., Campbell J. L., Kowalczykowski S. C., 2010 DNA end resection by Dna2-Sgs1-RPA and its stimulation by Top3-Rmi1 and Mre11-Rad50-Xrs2. 467: 112–116.

Cejka P., Plank J. L., Dombrowski C. C., Kowalczykowski S. C., 2012 Decatenation of DNA by the *S. cerevisiae* Sgs1-Top3-Rmi1 and RPA complex: a mechanism for disentangling chromosomes. Mol Cell 47: 886–896.

Chakraverty R. K., Kearsey J. M., Oakley T. J., Grenon M., La Torre Ruiz de M. A., Lowndes N. F., Hickson I. D., 2001 Topoisomerase III acts upstream of Rad53p in the S-phase DNA damage checkpoint. Mol Cell Biol 21: 7150–7162.

Chan K. L., Hickson I. D., 2011 New insights into the formation and resolution of ultra-fine anaphase bridges. Semin. Cell Dev. Biol. 22: 906–912.

Chan Y. W., Fugger K., West S. C., 2018 Unresolved recombination intermediates lead to ultra-fine anaphase bridges, chromosome breaks and aberrations. Nat Cell Biol 20: 92–103.

Chang M., Bellaoui M., Zhang C., Desai R., Morozov P., Delgado-Cruzata L., Rothstein R., Freyer G. A., Boone C., Brown G. W., 2005 *RMI1/NCE4*, a suppressor of genome instability, encodes a member of the RecQ helicase/Topo III complex. EMBO J 24: 2024–2033.

Chen S. Y., Tsubouchi T., Rockmill B., Sandler J. S., Richards D. R., Vader G., Hochwagen A., Roeder G. S., Fung J. C., 2008 Global analysis of the meiotic crossover landscape. Dev Cell 15: 401–415.

Cho R. J., Campbell M. J., Winzeler E. A., Steinmetz L., Conway A., Wodicka L., Wolfsberg T. G., Gabrielian A. E., Landsman D., Lockhart D. J., Davis R. W., 1998 A genome-wide transcriptional analysis of the mitotic cell cycle. Mol Cell 2: 65–73.

Chu S., Herskowitz I., 1998 Gametogenesis in yeast is regulated by a transcriptional cascade dependent on Ndt80. Mol Cell 1: 685–696.

Chu W. K., Hickson I. D., 2009 RecQ helicases: multifunctional genome caretakers. Nat. Rev. Cancer 9: 644–654.

Clyne R. K., Katis V. L., Jessop L., Benjamin K. R., Herskowitz I., Lichten M., Nasmyth K., 2003 Polo-like kinase Cdc5 promotes chiasmata formation and cosegregation of sister centromeres at meiosis I. Nat Cell Biol 5: 480–485.

Cohen-Fix O., Koshland D., 1997 The anaphase inhibitor of *Saccharomyces cerevisiae* Pds1p is a target of the DNA damage checkpoint pathway. Proc. Natl. Acad. Sci. U.S.A. 94: 14361–14366.

Cohen-Fix O., Peters J. M., Kirschner M. W., Koshland D., 1996 Anaphase initiation in *Saccharomyces cerevisiae* is controlled by the APC-dependent degradation of the anaphase inhibitor Pds1p. Genes Dev 10: 3081–3093.

Crickard J. B., Greene E. C., 2019 Helicase mechanisms during homologous recombination in *Saccharomyces cerevisiae*. Annu. Rev. Biophys. 48: pannurev–biophys–052118–115418.

Dayani Y., Simchen G., Lichten M., 2011 Meiotic recombination intermediates are resolved with minimal crossover formation during return-to-growth, an analogue of the mitotic cell cycle. PLoS Genet 7: e1002083.

De Muyt A., Jessop L., Kolar E., Sourirajan A., Chen J., Dayani Y., Lichten M., 2012 BLM helicase ortholog Sgs1 is a central regulator of meiotic recombination intermediate metabolism. Mol Cell 46: 43–53.

Ehmsen K. T., Heyer W.-D., 2008 Biochemistry of Meiotic Recombination: Formation, Processing, and Resolution of Recombination Intermediates. Genome Dynamics and Stability 3: 91–164.

Fabre F., Chan A., Heyer W.-D., Gangloff S., 2002 Alternate pathways involving Sgs1/Top3, Mus81/Mms4, and Srs2 prevent formation of toxic recombination intermediates from single-stranded gaps created by DNA replication. 99: 16887–16892.

Fasching C. L., Cejka P., Kowalczykowski S. C., Heyer W.-D., 2015 Top3-Rmi1 dissolve Rad51-mediated D loops by a topoisomerase-based mechanism. Mol Cell 57: 595–606.

Foiani M., Marini F., Gamba D., Lucchini G., Plevani P., 1994 The B subunit of the DNA polymerase alpha-primase complex in Saccharomyces cerevisiae executes an essential function at the initial stage of DNA replication. Mol Cell Biol 14: 923–933.

Friedlander G., Joseph-Strauss D., Carmi M., Zenvirth D., Simchen G., Barkai N., 2006 Modulation of the transcription regulatory program in yeast cells committed to sporulation. Genome Biol 7: R20.

Gangloff S., McDonald J. P., Bendixen C., Arthur L., Rothstein R., 1994 The yeast type I topoisomerase Top3 interacts with Sgs1, a DNA helicase homolog: a potential eukaryotic reverse gyrase. Mol Cell Biol 14: 8391–8398.

Giannattasio M., Zwicky K., Follonier C., Foiani M., Lopes M., Branzei D., 2014 Visualization of recombination-mediated damage bypass by template switching. Nat Struct Mol Biol.

Gloor G. B., Nassif N. A., Johnson-Schlitz D. M., Preston C. R., Engels W. R., 1991 Targeted gene replacement in *Drosophila* via P element-induced gap repair. Science 253: 1110–1117.

Goldfarb T., Lichten M., 2010 Frequent and efficient use of the sister chromatid for DNA double-strand break repair during budding yeast meiosis. PLoS Biol 8: e1000520.

Goyon C., Lichten M., 1993 Timing of molecular events in meiosis in *Saccharomyces cerevisiae*: stable heteroduplex DNA is formed late in meiotic prophase. Mol Cell Biol 13: 373–382.

Gravel S., Chapman J. R., Magill C., Jackson S. P., 2008 DNA helicases Sgs1 and BLM promote DNA double-strand break resection. Genes Dev 22: 2767–2772.

Harrison J. C., Haber J. E., 2006 Surviving the Breakup: The DNA Damage Checkpoint. Annu Rev Genet 40: 209–235.

Heijink A. M., Krajewska M., van Vugt M. A. T. M., 2013 The DNA damage response during mitosis. Mutat Res 750: 45–55.

Ira G., Malkova A., Liberi G., Foiani M., Haber J. E., 2003 Srs2 and Sgs1-Top3 suppress crossovers during double-strand break repair in yeast. Cell 115: 401–411.

Jessop L., Lichten M., 2008 Mus81/Mms4 endonuclease and Sgs1 helicase collaborate to ensure proper recombination intermediate metabolism during meiosis. Mol Cell 31: 313–323.

Jessop L., Allers T., Lichten M., 2005 Infrequent co-conversion of markers flanking a meiotic recombination initiation site in *Saccharomyces cerevisiae*. Genetics 169: 1353–1367.

Kane S. M., Roth R., 1974 Carbohydrate metabolism during ascospore development in yeast. J Bacteriol 118: 8–14.

Kaur H., Ahuja J. S., Lichten M., 2018 Methods for Controlled Protein Depletion to Study Protein Function during Meiosis. Meth Enzymol 601: 331–357.

Kaur H., De Muyt A., Lichten M., 2015 Top3-Rmi1 DNA single-strand decatenase is integral to the formation and resolution of meiotic recombination intermediates. Mol Cell 57: 583–594.

Kubota T., Nishimura K., Kanemaki M. T., Donaldson A. D., 2013 The Elg1 Replication Factor C-like Complex Functions in PCNA Unloading during DNA Replication. Mol Cell 50: 273–280.

Larsen N. B., Hickson I. D., 2013 RecQ Helicases: Conserved Guardians of Genomic Integrity. In:Spies M (Ed.), DNA Helicases and DNA Motor Proteins, DNA Helicases and DNA Motor Proteins. Springer New York, New York, NY, pp. 161–184.

Lynn A., Soucek R., Börner G. V., 2007 ZMM proteins during meiosis: crossover artists at work. Chromosome Res 15: 591–605.

Mancera E., Bourgon R., Brozzi A., Huber W., Steinmetz L. M., 2008 High-resolution mapping of meiotic crossovers and non-crossovers in yeast. 454: 479–485.

Mankouri H. W., Ashton T. M., Hickson I. D., 2011 Holliday junction-containing DNA structures persist in cells lacking Sgs1 or Top3 following exposure to DNA damage. Proc Natl Acad Sci USA 108: 4944–4949.

Martini E., Borde V., Legendre M., Audic S., Regnault B., Soubigou G., Dujon B., Llorente B., 2011 Genome-wide analysis of heteroduplex DNA in mismatch repair-deficient yeast cells reveals novel properties of meiotic recombination pathways. PLoS Genet 7: e1002305.

Matos J., West S. C., 2014 Holliday junction resolution: regulation in space and time. DNA Repair (Amst) 19: 176–181.

Matos J., Blanco M. G., West S. C., 2013 Cell-Cycle Kinases Coordinate the Resolution of Recombination Intermediates with Chromosome Segregation. Cell Reports 4: 76–86.

Matos J., Blanco M. G., Maslen S., Skehel J. M., West S. C., 2011 Regulatory control of the resolution of DNA recombination intermediates during meiosis and mitosis. Cell 147: 158–172.

Mimitou E. P., Symington L. S., 2009 DNA end resection: many nucleases make light work. DNA Repair (Amst) 8: 983–995.

Morawska M., Ulrich H. D., 2013 An expanded tool kit for the auxin-inducible degron system in budding yeast. Yeast 30: 341–351.

Mullen J. R., Kaliraman V., Ibrahim S. S., Brill S. J., 2001 Requirement for three novel protein complexes in the absence of the Sgs1 DNA helicase in *Saccharomyces cerevisiae*. Genetics 157: 103–118.

Mullen J. R., Nallaseth F. S., Lan Y. Q., Slagle C. E., Brill S. J., 2005 Yeast Rmi1/Nce4 controls genome stability as a subunit of the Sgs1-Top3 complex. Mol Cell Biol 25: 4476–4487.

Nishimura K., Fukagawa T., Takisawa H., Kakimoto T., Kanemaki M., 2009 An auxin-based degron system for the rapid depletion of proteins in nonplant cells. Nat Methods 6: 917–922.

Nyberg K. A., Michelson R. J., Putnam C. W., Weinert T. A., 2002 Toward maintaining the genome: DNA damage and replication checkpoints. Annu Rev Genet 36: 617–656.

Oakley T. J., Goodwin A., Chakraverty R. K., Hickson I. D., 2002 Inactivation of homologous recombination suppresses defects in topoisomerase III-deficient mutants. DNA Repair (Amst.) 1: 463–482.

Oh S. D., Jessop L., Lao J. P., Allers T., Lichten M., Hunter N., 2009 Stabilization and electrophoretic analysis of meiotic recombination intermediates in *Saccharomyces cerevisiae*. Methods Mol Biol 557: 209–234.

Oh S. D., Lao J. P., Hwang P. Y.-H., Taylor A. F., Smith G. R., Hunter N., 2007 BLM ortholog, Sgs1, prevents aberrant crossing-over by suppressing formation of multichromatid joint molecules. Cell 130: 259–272.

Oh S. D., Lao J. P., Taylor A. F., Smith G. R., Hunter N., 2008 RecQ helicase, Sgs1, and XPF family endonuclease, Mus81-Mms4, resolve aberrant joint molecules during meiotic recombination. Mol Cell 31: 324–336.

Piazza A., Heyer W.-D., 2019 Moving forward one step back at a time: reversibility during homologous recombination. Curr Genet.

Piazza A., Shah S. S., Wright W. D., Gore S. K., Koszul R., Heyer W.-D., 2019 Dynamic processing of displacement loops during recombinational DNA repair. Mol Cell 73: 1255–1266.e4.

Plank J. L., Wu J., Hsieh T.-S., 2006 Topoisomerase IIIalpha and Bloom’s helicase can resolve a mobile double Holliday junction substrate through convergent branch migration. 103: 11118–11123.

Pyatnitskaya A., Borde V., De Muyt A., 2019 Crossing and zipping: molecular duties of the ZMM proteins in meiosis. Chromosoma 4: e1000042–18.

Schiller C. B., Seifert F. U., Linke-Winnebeck C., Hopfner K.-P., 2014 Structural studies of DNA end detection and resection in homologous recombination. Cold Spring Harb Perspect Biol 6: a017962–a017962.

Shor E., Gangloff S., Wagner M., Weinstein J., Price G., Rothstein R., 2002 Mutations in homologous recombination genes rescue *top3* slow growth in *Saccharomyces cerevisiae*. Genetics 162: 647–662.

Shor E., Weinstein J., Rothstein R., 2005 A genetic screen for *top3* suppressors in *Saccharomyces cerevisiae* identifies *SHU1, SHU2, PSY3 and CSM2*: four genes involved in error-free DNA repair. Genetics 169: 1275–1289.

Simchen G., 2009 Commitment to meiosis: what determines the mode of division in budding yeast? Bioessays 31: 169–177.

Sourirajan A., Lichten M., 2008 Polo-like kinase Cdc5 drives exit from pachytene during budding yeast meiosis. Genes Dev 22: 2627–2632.

Subramanian V. V., Hochwagen A., 2014 The Meiotic Checkpoint Network: Step-by-Step through Meiotic Prophase. Cold Spring Harb Perspect Biol 6.

Szostak J. W., Orr-Weaver T. L., Rothstein R. J., Stahl F. W., 1983 The double-strand-break repair model for recombination. Cell 33: 25–35.

Tang S., Wu M. K. Y., Zhang R., Hunter N., 2015 Pervasive and essential roles of the Top3-Rmi1 decatenase orchestrate recombination and facilitate ehromosome segregation in meiosis. Mol Cell 57: 607–621.

Tsubouchi H., Argunhan B., Tsubouchi T., 2018 Exiting prophase I: no clear boundary. Curr Genet 64: 423–427.

van Brabant A. J., Ye T., Sanz M., German J. L. III, Ellis N. A., Holloman W. K., 2000 Binding and melting of D-loops by the Bloom syndrome helicase. Biochemistry 39: 14617–14625.

Visintin C., Tomson B. N., Rahal R., Paulson J., Cohen M., Taunton J., Amon A., Visintin R., 2008 APC/C-Cdh1-mediated degradation of the Polo kinase Cdc5 promotes the return of Cdc14 into the nucleolus. Genes Dev 22: 79–90.

Wallis J. W., Chrebet G., Brodsky G., Rolfe M., Rothstein R., 1989 A hyper-recombination mutation in *S. cerevisiae* identifies a novel eukaryotic topoisomerase. Cell 58: 409–419.

Wells W. A., 1996 The spindle-assembly checkpoint: aiming for a perfect mitosis, every time. Trends Cell Biol 6: 228–234.

Winey M., Bloom K., 2012 Mitotic spindle form and function. Genetics 190: 1197–1224.

Winter E., 2012 The Sum1/Ndt80 transcriptional switch and commitment to meiosis in *Saccharomyces cerevisiae*. Microbiology and Molecular Biology Reviews 76: 1–15.

Wu L., Hickson I. D., 2003 The Bloom’s syndrome helicase suppresses crossing over during homologous recombination. Nature 426: 870–874.

Wu L., Bachrati C. Z., Ou J., Xu C., Yin J., Chang M., Wang W., Li L., Brown G. W., Hickson I. D., 2006 BLAP75/RMI1 promotes the BLM-dependent dissolution of homologous recombination intermediates. Proc Natl Acad Sci USA 103: 4068–4073.

Wyatt H. D. M., West S. C., 2014 Holliday junction resolvases. Cold Spring Harb Perspect Biol 6: a023192–a023192.

Xaver M., Huang L., Chen D., Klein F., 2013 Smc5/6-Mms21 prevents and eliminates inappropriate recombination intermediates in meiosis. PLoS Genet 9: e1004067.

Zakharyevich K., Ma Y., Tang S., Hwang P. Y.-H., Boiteux S., Hunter N., 2010 Temporally and biochemically distinct activities of Exo1 during meiosis: double-strand break resection and resolution of double Holliday junctions. Mol Cell 40: 1001–1015.

Zakharyevich K., Tang S., Ma Y., Hunter N., 2012 Delineation of joint molecule resolution pathways in meiosis identifies a crossover-specific resolvase. Cell 149: 334–347.

Zenvirth D., Loidl J., Klein S., Arbel A., Shemesh R., Simchen G., 1997 Switching yeast from meiosis to mitosis: double-strand break repair, recombination and synaptonemal complex. Genes Cells 2: 487–498.

Zhu Z., Chung W.-H., Shim E. Y., Lee S. E., Ira G., 2008 Sgs1 helicase and two nucleases Dna2 and Exo1 resect DNA double-strand break ends. Cell 134: 981–994.

